# Therapeutic Prime/Pull Vaccination of HSV-2 Infected Guinea Pigs with the Ribonucleotide Reductase 2 (RR2) Protein and CXCL11 Chemokine Boosts Antiviral Local Tissue-Resident and Effector Memory CD4^+^ and CD8^+^ T Cells and Protects Against Recurrent Genital Herpes

**DOI:** 10.1101/2023.08.08.552454

**Authors:** Afshana Quadiri, Swayam Prakash, Nisha Rajeswari Dhanushkodi, Mahmoud Singer, Latifa Zayou, Amin Mohammed Shaik, Miyo Sun, Berfin Suzer, Lauren Lau, Amruth Chilukurri, Hawa Vahed, Hubert Schaefer, Lbachir BenMohamed

## Abstract

Following acute herpes simplex virus type 2 (HSV-2) infection, the virus undergoes latency in sensory neurons of the dorsal root ganglia (DRG). Intermittent virus reactivation from latency and shedding in the vaginal mucosa (VM) causes recurrent genital herpes. While T-cells appear to play a role in controlling virus reactivation and reducing the severity of recurrent genital herpes, the mechanisms for recruiting these T-cells into DRG and VM tissues remain to be fully elucidated. The present study investigates the effect of CXCL9, CXCL10, and CXCL11 T-cell-attracting chemokines on the frequency and function of DRG- and VM-resident CD4^+^ and CD8^+^ T cells and its effect on the frequency and severity of recurrent genital herpes. HSV-2 latent-infected guinea pigs were immunized intramuscularly with the HSV-1 RR2 protein *(Prime)* and subsequently treated intravaginally with the neurotropic adeno-associated virus type 8 (AAV-8) expressing CXCL9, CXCL10, or CXCL11 T-cell-attracting chemokines (*Pull)*. Compared to the RR2 therapeutic vaccine alone, the RR2/CXCL11 prime/pull therapeutic vaccine significantly increased the frequencies of functional tissue-resident (T_RM_ cells) and effector (T_EM_ cells) memory CD4^+^ and CD8^+^ T cells in both DRG and VM tissues. This was associated with less virus shedding in the healed genital mucosal epithelium and reduced frequency and severity of recurrent genital herpes. These findings confirm the role of local DRG- and VM-resident CD4^+^ and CD8^+^ T_RM_ and T_EM_ cells in reducing virus reactivation shedding and the severity of recurrent genital herpes and propose the novel prime/pull vaccine strategy to protect against recurrent genital herpes.

**IMPORTANCE:** The present study investigates a novel prime/pull therapeutic vaccine strategy to protect against recurrent genital herpes in the guinea pig model. HSV-2 infected guinea pigs were vaccinated using a recombinantly expressed herpes tegument protein-RR2 (*prime*), followed by intravaginal treatment with the neurotropic adeno-associated virus type 8 (AAV-8) expressing CXCL9, CXCL10, or CXCL11 T-cell-attracting chemokines (pull). The RR2/CXCL11 prime/pull therapeutic vaccine elicited a significant reduction in virus shedding in the vaginal mucosa and decreased the severity and frequency of recurrent genital herpes. This protection was associated with increased frequencies of functional tissue-resident (T_RM_ cells) and effector (T_EM_ cells) memory CD4^+^ and CD8^+^ T cells infiltrating latently infected DRG tissues and the healed regions of the vaginal mucosa. These findings shed light on the role of tissue-resident (T_RM_ cells) and effector (T_EM_ cells) memory CD4^+^ and CD8^+^ T cells in DRG and vaginal mucosa (VM) tissues in protection against recurrent genital herpes and propose the prime/pull therapeutic vaccine strategy in combating genital herpes.

**TWEET:** A therapeutic RR2/CXCL11 prime/pull vaccine reduced recurrent genital herpes more effectively than therapeutic vaccination with a subunit HSV RR2 antigen alone.

## INTRODUCTION

Herpes simplex virus type 2 (HSV-2) affects both women and men (1, 2), however, women are more susceptible to the infection. Approximately 315 million women aged 5-49 years old are currently infected globally (1, 2). After exposure of vaginal mucosa (VM) to HSV-2, the virus replicates in the mucosal epithelial cells leading to the development of acute genital herpetic lesions (2-5). Once the acute primary infection is cleared, the virus establishes a lifelong latent infection. The virus enters the nerve termini innervating peripheral vaginal tissues and is subsequently transported by retrograde to the nucleus of the sensory neurons of dorsal root ganglia (DRG) where it establishes a dormant state within the neuronal cells (6, 7).

More than 80% of HSV-2-seropositive women are unaware of their infection as they never develop any apparent recurrent symptoms (1). In contrast, the symptomatic women often display sporadic reactivation, leading to recurrent genital lesions and painful blisters that can burst and form ulcers (6). The virus occasionally reactivates and sheds asymptomatically, even without visible lesions. Such women, being unaware, can contribute significantly to the transmission of the virus, emphasizing the need for an antiviral therapeutic vaccine to prevent or reduce virus reactivation and/or its shedding in the genital tract. In addition, neonatal infections can be severe and result in serious diseases with high rates of morbidity and mortality. Despite the availability of many intervention strategies, such as sexual behavior education, barrier methods, and antiviral drug therapies (e.g., Acyclovir and derivatives), eliminating or at least reducing recurrent genital herpes remains a challenge (2, 8-11). Besides, antiviral drugs, such as Acyclovir, can neither prevent de novo infection (initial infection) nor clear the virus completely. They work primarily in reducing the severity and duration of symptoms by inhibiting viral replication during active outbreaks. An effective antiviral therapeutic vaccine may serve as the best approach to protect from recurrent genital herpetic disease (4, 12). An antiviral therapeutic vaccine stimulates the immune system to target and control an existing infection. Ideally, a therapeutic vaccine for genital herpes would help reduce virus reactivation, limit shedding, and potentially decrease the frequency and severity of recurrent outbreaks.

The acquired immune responses that develop following exposure to the virus are not sufficient for protection against recurrent genital herpes (13-15). More recently, studies are investigating a successful therapeutic vaccine that can boost immune responses stronger and/or different than the acquired immunity induced by the virus (16-19). In 1988, a study by Stanberry et al. provided early evidence that a vaccine could reduce the number of recurrent genital herpes outbreaks using the guinea pig model of genital herpes. This study demonstrated the efficacy of immunization in reducing the frequency of recurrences. In recent years, several sub-clinical experiments have supported these preclinical findings using protein-based subunit vaccines, often administered with a potent adjuvant (a substance that enhances the immune response). Vaccination with these proteins has been found to reduce the rates of recurrent lesions (visible outbreaks) or recurrent shedding by approximately 50%.

Interestingly, over the last two decades, only a single subunit protein vaccine strategy, based on the HSV-2 glycoprotein D (gD), delivered with or without gB, has been tested and retested in clinical trials (18, 20). Despite inducing strong neutralizing antibodies; this subunit vaccine strategy proved unsuccessful in clinical trials (21). Previous studies have identified other antigenic tegument proteins by screening HSV-2 open-reading frames (ORFs) with antibodies and T cells from HSV-2 seropositive individuals (22). However, aside from three reports, first by our group in 2012 (23, 24) and later by Genocea Biosciences, Inc. in 2014 (25), comparison of the repertoire of HSV-2 proteins, encoded by the over 84 open reading frames- (ORF-) of the HSV-2 152-kb genome, recognized by antibodies and T cells from HSV-2 seropositive symptomatic versus asymptomatic individuals is largely unknown. Previously, we demonstrated that the HSV-2 specific RR2 protein-based subunit therapeutic vaccine elicited a significant reduction in virus shedding and decreased the severity and frequency of recurrent genital herpes lesions (26). Our lab previously determined that the RR2 protein is frequently and highly recognized by antibodies and T cells in naturally “protected” asymptomatic individuals. Additionally, the protein boosted the number and function of antiviral tissue-resident memory CD4^+^ and CD8^+^T_RM_ cells, locally within the DRG and vaginal mucocutaneous tissues, leading to better protection against recurrent herpes (26).

In the present study, we sought to improve protein-based vaccination by combining it with a prime and pull strategy. The strategy involves conventional parenteral vaccination using HSV-2 specific RR2 protein to elicit systemic T-cell responses (prime), followed by recruitment of activated T cells via administration of an adenovirus expressing chemokine or T cell attractant (pull), for those T cells to establish long-term protective immunity. The adeno-associated virus type 8 (AAV8) was used to express the chemokines as it efficiently targets the enteric nervous system in guinea pigs. The immunization results showed that the primary CD8^+^T cell responses were of similar magnitudes in the spleen, whereas the frequency and number of CD8^+^T cells in the vaginal mucocutaneous tissue were significantly higher in vaccinated mice treated with adenovirus-expressing chemokine as compared with the control immunized mice without the “pull”. Furthermore, the action of the chemokine pull was not restricted to the genital mucosa, as CD8^+^ T cell recruitment was also observed in the DRG. Importantly, the prime and pull strategy conferred near complete protection against the primary challenge of genital HSV-2 infection compared with the prime alone. In this study, we extend this application to therapeutic vaccines and demonstrate that the frequency of recurrent disease and recurrent vaginal shedding is reduced most effectively by the combination of prime (protein vaccine) and pull (Adenovirus expressing chemokine).

## MATERIALS AND METHODS

### Animals

Female guinea pigs (*Hartley strain*, Charles River Laboratories, San Diego, CA) weighing 275-350 g (5-6 weeks old) were housed at the University of California, Irvine vivarium. The Institutional Animal Care and Use Committee of the University of California, Irvine, reviewed and approved the protocol for these studies (IACUC # AUP-22-086). A group size of 10 had 90% power to detect a difference of two-fold or higher between experimental group means with a significance level of 0.05.

### Vaccine candidate

We used recombinantly expressed ‘RR2’ protein antigens from HSV2 as RR2 is highly recognized by T cells from naturally protected” asymptomatic individuals.

### Infection and Immunization of Guinea Pigs

Throughout this study, we used the MS strain of HSV-2, generously gifted by Dr. David Bernstein (Cincinnati Children’s Hospital Medical Center, University of Cincinnati, OH). Guinea pigs (*n* = 30) were infected intravaginally with 5 x 10^5^ pfu of HSV-2 (strain MS). Once the acute infection was resolved, latently infected animals were vaccinated intramuscularly twice in the right hind calf muscle on day 15 and day 25 post-infection. Animals were immunized on day 15 with 20 μg and on day 25 with 10 μg of RR2 protein mixed with 100 μg CpG/guinea pig and 150μg alum. Animals were mock-immunized with the CpG oligonucleotide (5’-TCGTCGTTGTCGTTTTGTCGTT-3’) (Trilink Inc., Santa Fe Springs, CA) using 100 μg CpG/guinea pig and 150μg alum (Alhydrogel, Accurate Chemical and Scientific Corp., Westbury, NY).

### Monitoring of primary or recurrent HSV-2 disease in guinea pigs

Guinea pigs were examined for vaginal lesions and were recorded for each animal daily starting right after the second immunization on a scale of 0 to 4, where 0 reflects no disease, 1 reflects redness, 2 reflects a single lesion, 3 reflects coalesced lesions, and 4 reflects ulcerated lesions.

### Bulk RNA sequencing on sorted CD8+ T cells

RNA was isolated from the sorted CD8+ T cells using the Direct-zol RNA MiniPrep (Zymo Research, Irvine, CA) according to the manufacturer’s instructions. RNA concentration and integrity were determined using the Agilent 2100 Bioanalyzer. Sequencing libraries were constructed using TruSeq Stranded Total RNA Sample Preparation Kit (Illumina, San Diego, CA). Briefly, rRNA was first depleted using the RiboGone rRNA removal kit (Clonetech Laboratories, Mountain View, CA) before the RNA was fragmented, converted to double-stranded cDNA and ligated to adapters, amplified by PCR and selected by size exclusion. Following quality control for size, quality, and concentrations, libraries were multiplexed and sequenced to single-end 100-bp sequencing using the Illumina HiSeq 4000 platform.

### Differential gene expression analysis

Differentially expressed genes (DEGs) were analyzed using Integrated Differential Expression and Pathway analysis tools. These system tools seamlessly connect 63 R/Bioconductor packages, two web services, and comprehensive annotation and pathway databases for guinea pigs. The expression matrix of DEGs was filtered and converted to Ensembl gene identifiers, and the preprocessed data were used for exploratory data analysis, including k-means clustering and hierarchical clustering. The pairwise comparison of immunized and non-immunized guinea pigs was performed using the DESeq2 package with a single-Andover rate threshold (<0.5. and fold change and >1.5).

Moreover, a hierarchical clustering tree and network of enriched GO/KEGG terms were constructed to visualize the potential relationship. Gene Set Enrichment Analysis (GSEA) method was performed to investigate the related signal pathways activated among protective and non-protective groups. The parametric Gene Set Enrichment Analysis (PSGEA) method was applied based on data curated in Gene Ontology and KEGG. Pathway significance cutoff with a false discovery date (FDR) ≥ 0.2 was used.

### Real-time qPCR for HSV-2 Quantification from Vaginal Swabs and dorsal root ganglia

Vaginal swabs were collected daily using a Dacron swab (type 1; Spectrum Laboratories, Los Angeles, CA) starting from day 35 until day 65 post-challenge. Individual swabs were transferred to a 2 mL sterile cryogenic vial containing 1ml culture medium and stored at -80°C until use. On day 65 post-challenge, twelve lower lumbar and sacral dorsal root ganglia (DRG) per guinea pig were collected by cutting through the lumbar end of the spine. DNA was isolated from the collected vaginal swab and DRG of guinea pigs by using DNeasy blood and tissue kits (Qiagen). The presence of HSV-2 DNA was quantified by real-time PCR (StepOnePlus Real-Time PCR System) with 50-100 ng vaginal swab DNA or 250 ng of DRG DNA. HSV-2 DNA copy number was determined using purified HSV-2 DNA (Advanced Biotechnologies, Columbia, MD) and based on a standard curve of the *C_T_* values that were generated with 50,000, 5,000, 500, 50, and 5 copies of DNA and run in triplicates, samples analyzed in duplicates. Samples with <150 copies/ml by 40 cycles or only positive in one of two wells were reported as negative. Primer and probe sequences for HSV-2 Us9 were: primer forward, 5′-GGCAGAAGCCTACTACTCGGAAA-3′, and reverse 5′-CCATGCGCACGAGGAAGT-3′, and probe with reporter dye 5′-FAM-CGAGGCCGCCAAC-MGBNFQ-3′ (FAM, 6-carboxyfluorescein). All reactions were performed using TaqMan gene expression master mix (Applied Biosystems), and data were collected and analyzed on StepOnePlus real-time PCR system.

### Splenocyte isolation

Spleens were harvested from guinea pigs at 80 days post-infection. Spleens were placed in 10 ml of cold PBS with 10% fetal bovine serum (FBS) and 2X antibiotic– antimycotic (Life Technologies, Carlsbad, CA). Spleens were minced finely and sequentially passed through a 100 µm mesh and a 70 µm mesh (BD Biosciences, San Jose, CA). Cells were then pelleted via centrifugation at 400 × *g* for 10 minutes at 4°C. Red blood cells were lysed using a lysis buffer and washed again. Isolated splenocytes were diluted to 1 × 10^6^ viable cells per ml in RPMI media with 10% (v/v) FBS and 2 × antibiotic–antimycotic. Viability was determined by Trypan blue staining.

### Isolation of lymphocytes from the guinea pig’s vaginal mucosa

Vaginal mucosa was removed from the guinea pigs. The genital tract was minced into fine pieces and transferred into a new tube with fresh RPMI-10 containing collagenase and digested at 37 °C for two hours on a rocker set to vigorously. The digested tissue suspension was then passed through a 100 μm cell strainer on ice, followed by centrifugation. Lymphocytes in the cell pellets were separated using Percoll gradients by centrifugation at 900 x g, at room temperature, for 20 minutes with the brake-off. The lymphocytes at the interface layer between 40% and 70% Percoll layers were harvested, washed with RPMI 1:3, and spun down at 740 x g.

### Flow cytometry analysis

Vaginal mucosa cells and splenocytes were analyzed by flow cytometry using the following antibodies: mouse anti-guinea pig CD8 (clone MCA752F, Bio-Rad Laboratories, Hercules, CA), mouse anti-guinea pig CD4 (clone MCA749PE, Bio-Rad Laboratories), anti-mouse CRTAM (clone 11-5, Biolegend, San Diego, CA), hamster anti-mouse PD-1 clone J43, BD Biosciences, San Jose, CA), anti-mouse/human CD44 (clone IM7, Biolegend), anti-mouse CD69 (clone H1.2F3, BD Biosciences, San Jose, CA), anti-mouse CXCR3 (clone CXCR3-173, Biolegend) and anti-mouse CD103 (clone 2E7, Biolegend). For surface staining, mAbs against various cell markers were added to a total of 1X10^6^ cells in phosphate-buffered saline containing 1% FBS and 0.1% sodium azide (fluorescence-activated cell sorter [FACS] buffer) and left for 45 minutes at 4°C. At the end of the incubation period, the cells were washed twice with FACS buffer. A total of 100,000 events were acquired by the LSRII (Becton Dickinson, Mountain View, CA), followed by analysis using FlowJo software (TreeStar, Ashland, OR).

### Statistical analyses

Data for each assay were compared by analysis of variance (ANOVA) and Student’s t-test using GraphPad Prism version 5 (La Jolla, CA). Differences between the groups were identified by ANOVA and multiple comparison procedures. Data are expressed as the mean ± SD. Results were considered statistically significant at *P* < 0.05.

## RESULTS

**1. *CXCL11/CXCR3 axis for T cell activation: A target for novel “prime-pull” therapeutic herpes vaccine:*** Guinea pigs (*n* = 10) were infected intravaginally with 5 x 10^5^ pfu of HSV-2 (strain MS). Once the acute infection was resolved, latently infected animals were vaccinated intramuscularly twice, on days 15 and 25 post-infection, with individual HSV-2 antigen RR2 emulsified in Alum + CpG adjuvants. Mock-vaccinated guinea pigs, which received Alum + CpG adjuvants alone, were used as negative control (Mock). Using bulk RNA sequencing, we detected differential expression of 1,243 (upregulated = 724, downregulated = 519) genes in tissue-resident CD8^+^T cells FACS-sorted from vaginal mucosa of HSV-2 immunized and non-immunized guinea pigs (**Figs. 1A** and **1B**, *left panel*). With gene set enrichment analysis on these differentially expressed genes, we found T cell activation (*P* = 0.02, logFC = 4.12), T cell proliferation (P = 0.02, logFC =3.68), TCR signaling (*P* = 0.01, logFC = 2.15), and Cytokine secretion (P = 0.01, logFC = 2.01) pathways to be upregulated in immunized group of guinea pigs (**Fig. 1B***, middle panel*). We detected a significant up-regulation of the gene for T cells attracting chemokine receptors in HSV-specific CD8^+^T_RM_ cells from an immunized group of guinea pigs suggestive of a heightened activation of T cell-attracting chemokines receptors may lead to increased infiltration/retention of protective antiviral T_RM_ cells observed in the VM of the immunized group of guinea pigs. The gene found to be significantly upregulated among immunized guinea pigs was CXCR3 (P = 0.004, logFC = 3.46). ((**Figs. 1D** and **1E**), which is a receptor for CXCL9, CXCL10 and CXCL11. These results suggest that CXCL9/10/11-CXCR3 activates tissue resident CD8^+^ T cell responses locally in the genital tissues.

**Figure 1:**
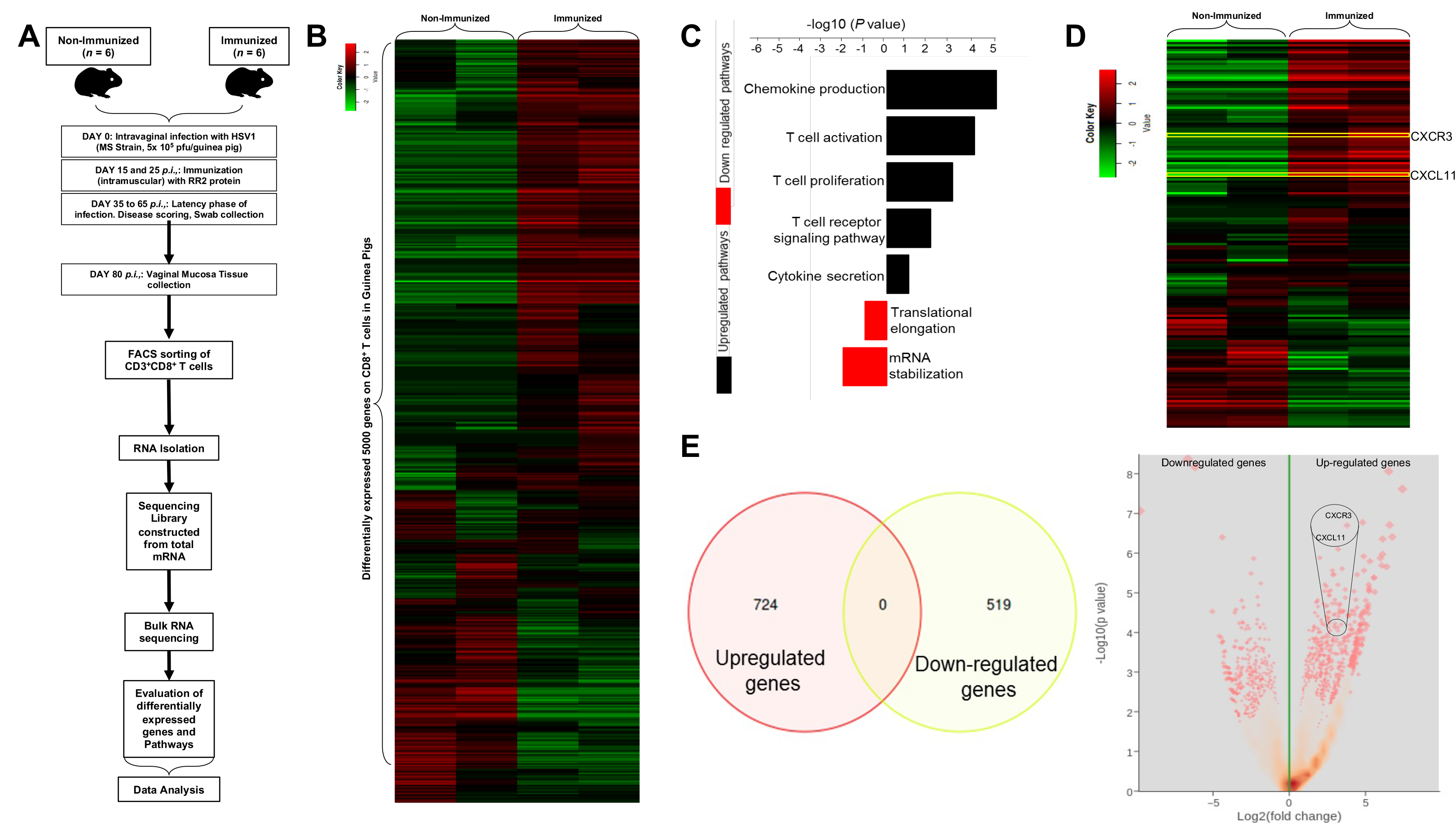
*Higher frequency of tissue-resident CD8^+^ T cells in healed vaginal mucosal cells of protected guinea pigs is associated with stimulation of the CXCL11/CXCR3 axis:* (**A**) Guinea pigs (*n* = 12) were infected intravaginally with 5 x 10^5^ pfu of HSV-2 (strain MS). Once the acute infection was resolved, latently infected animals were vaccinated intramuscularly twice, on days 15 and 25 post-infection with individual HSV-2 antigen RR2 emulsified in Alum + CpG adjuvants,Mock-vaccinated guinea pigs, which received Alum + CpG adjuvants alone, were used as negative control (*Mock*) (**B**) Heatmap showing differential gene expression (DGE) analysis in guinea pig CD8^+^ T cells. (**C**) GSEA Analysis showing differentially expressed immunological pathways in CD8^+^ T cells associated with HSV-2 infection in guinea pigs (**D**) GSEA analysis showing differentially upregulated Chemokine production pathway in protected guinea pigs. Upregulation of CXCR3, and its corresponding ligands CXCL9, CXCL10, and CXCL11 observed among guinea pigs infected with HSV-2 in the data obtained from bulk RNA sequencing (**E**) *Left panel:* Venn diagram showing the distribution of significantly differentially expressed genes among immunized and non-immunized guinea pigs, *Right panel:* Volcano plot showing upregulated m-RNA fold change expression for CXCR3, CXCL9, CXCL10, and CXCL11 among infected guinea pigs.

**2. *Therapeutic immunization of HSV-2 infected guinea pigs with vaccine candidate RR2 and treatment with adenovirus containing CXCL11 protects better against recurrent genital disease:*** Guinea pigs (n = 30) were infected intravaginally with 5 x 10^5^ pfu of HSV-2 (strain MS) (Fig. 2A). Once acute infection was resolved, latently infected animals were randomly divided into five groups (n = 6) and then vaccinated twice intramuscularly on days 15 and 25 post-infection with HSV-2 protein RR2 emulsified in Alum + CpG adjuvants. One week later, three groups were treated with an adenovirus containing CXCL9, CXCL10, and CXCL11 separately, while the other group remained untreated. Mock-vaccinated guinea pigs (*n* = 6), who received alum plus CpG adjuvants alone, were used as negative controls. Starting on day 35 until day 65, the guinea pigs were observed and scored regularly for genital lesions. RR2-vaccinated animals treated with chemokine and RR2 alone vaccinated animals exhibited significantly lower cumulative vaginal lesions (**Fig. 2C**) and an overall significant reduction in cumulative positive days of recurrence compared to the mock-vaccinated controls (**Fig. 2D**). RR2-vaccinated animals treated with CXCL11 displayed lowest cumulative vaginal lesions and an overall reduced cumulative positive days of recurrence compared to the other vaccinated and chemokine-treated animals (**Figs. 2C** and **2D**).

**Figure 2:**
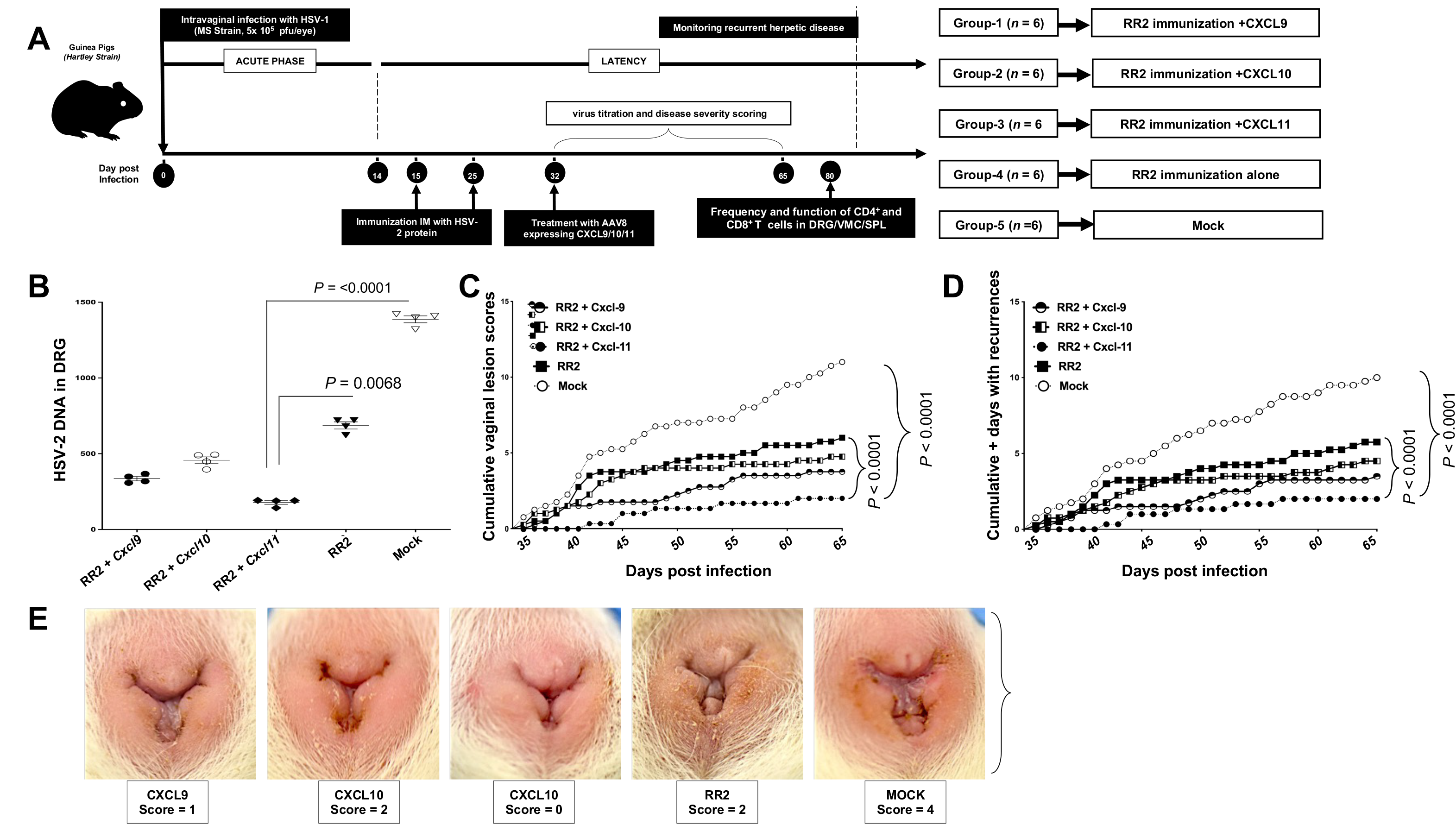
Protection against recurrent genital herpes infection and disease in HSV-2 infected guinea pigs following therapeutic prime/pull vaccination with the RR2 protein and adenovirus expressing chemokine CXCL9/10/11: (**A**) Timeline of HSV-2 infection, immunization, immunological and virological analyses. Guinea pigs (*n* = 30) were infected intravaginally on day 0 with 5 x 10 pfu of HSV-2 (strain MS). Once acute infection was resolved, the remaining latently infected animals were randomly divided into 5 groups (*n* = 6) and then vaccinated intramuscularly twice, on day 15 and day 25 post-infection, with 10 µg of the HSV-2 protein-based subunit vaccine-RR2 emulsified in Alum + CpG adjuvants. One Mock-vaccinated guinea pig that received Alum + CpG adjuvants alone was used as negative control (*Mock*). One week later, on day 32, one group was treated with an adenovirus expressing CXCL9, 2^nd^ group was treated with an adenovirus expressing CXCL10, 3^rd^ group was treated with an adenovirus expressing CXCL11, while the other group remained untreated (RR2 alone). From day 35 to day 80 post-infection (i.e. day 10 to day 55 after final immunization), vaccinated and non-vaccinated animals were observed daily for (*i*) the severity of genital herpetic lesions scored on a scale of 0 to 4, and pictures of genital areas taken; and (*ii*) vaginal swabs were collected daily from day 35 to day 65 post-infection (i.e. day 10 to day 40 after final immunization) to detect virus shedding and to quantify HSV-2 DNA copy numbers. (**B**) HSV-2 DNA copy numbers detected in the DRG of each vaccinated and mock-vaccinated guinea pig group. (**C**) Cumulative scoring of vaginal lesions observed during recurrent infection (virus titers). (**D**) Cumulative positive days with recurrent genital lesions. The severity of genital herpetic lesions was scored on a scale of 0 to 4, where 0 reflects no disease, 1 reflects redness, 2 reflects a single lesion, 3 reflects coalesced lesions, and 4 reflects ulcerated lesions. (**E**) Representative images of genital lesions in guinea pigs vaccinated with (*i*) RR2 and CXCL9 (*ii*) RR2 and CXCL10 (*iii*) RR2 and CXCL11 (*iv*) RR2 alone. The indicated *P* values show statistical significance between the HSV-2 vaccinated and mock-vaccinated control groups.

On day 80 post-infection, the RR2/Chemokine treated and RR2 alone vaccinated guinea pigs exhibited lower HSV-2 DNA copy numbers in the DRG than did mock-vaccinated controls (**Fig. 2B**), which was associated with a significant reduction in cumulative virus vaginal shedding in the vaccinated group as compared to the mock vaccinated control group. The severity of genital herpetic lesions scored on a scale of 0 to 4, also confirmed the cure of recurrent disease in RR2/CXCL11 treated and RR2 alone vaccinated guinea pigs (**Fig. 2E**). The lowest genital lesions were observed in guinea pigs vaccinated with RR2 protein and treated with CXCL11 (**Fig. 2E**, *middle panel*). RR2 + chemokine and RR2 alone vaccinated group was moderately protective against genital lesions (**Fig. 2E**). However, the mock vaccinated group showed no significant protection against recurrent genital herpes lesions (**Fig. 2E**, *right panel*). Altogether, these results indicate that therapeutic immunization with RR2 and treatment with AV containing CXCL11 protected HSV-2-seropositive guinea pigs against recurrent genital herpes infection and disease.

**3. *Therapeutic prime/pull vaccination of HSV-2 infected guinea pigs with RR2 protein/CXCL11 increased the frequencies of tissue-resident CD4^+^ and CD8^+^ T cells:*** Subsequently, we determined the frequencies of CD4^+^ and CD8^+^ T cells in the spleen (SPL), vaginal mucosa (VM), and dorsal root ganglia (DRG). Guinea pigs were infected and immunized, as detailed above. On day 80, after the second and final immunization treatment with AV containing CXCL9/10/11, vaccinated alone and control animals were euthanized, and the frequencies of SPL, DRG, and VM tissue-resident CD4^+^ and CD8^+^ T cells were detected by fluorescence-activated cell sorting (FACS). There were no significant differences observed in the frequencies of CD4^+^ and CD8^+^ T cells in the SPL of vaccinated guinea pigs compared to the mock vaccinated group. However, significantly higher frequencies of CD4^+^ and CD8^+^ T cells were induced in the RR2 vaccinated group following treatment with the chemokine, especially CXCL11, compared to RR2 vaccinated group alone and compared to those with the mock vaccinated group (i.e., adjuvant alone) in the VM and DRG tissues (Supplemental **Figs. S1A** and **S1B**).

**4. *Therapeutic prime/pull vaccination of HSV-2 infected guinea pigs with RR2 protein followed by CXCL11 treatment protected by bolstering effector and CXCR3 responses:*** We next determined the expression of CXCR3 expression on the CD8^+^ T cells. On day 80, after the second and final therapeutic prime/pull vaccination and treatment with chemokines, guinea pigs were euthanized, and single-cell suspensions from the SPL, VM, and DRG tissue were obtained, and the effector function of SPL, VM-resident, and DRG-resident CD8^+^ T cells was analyzed by observing the expression of CXCR3 on CD8^+^ T cells by FACS analysis. A non-significant difference was observed in the frequencies of CXCR3^+^ CD8^+^ T cells in the SPL of guinea pigs vaccinated with RR2 and treated with chemokines and RR2 protein alone compared to the mock vaccinated group (**Fig. 3C**). A higher frequency of CXCR3^+^CD8^+^ T cells was induced by the RR2 vaccinated group treated with chemokine especially CXCL11, followed by vaccination with the RR2 alone compared to those with the mock vaccinated group (i.e., adjuvant alone) in the VM and DRG tissues (**Fig. 3B** and **3C**).

**Figure 3:**
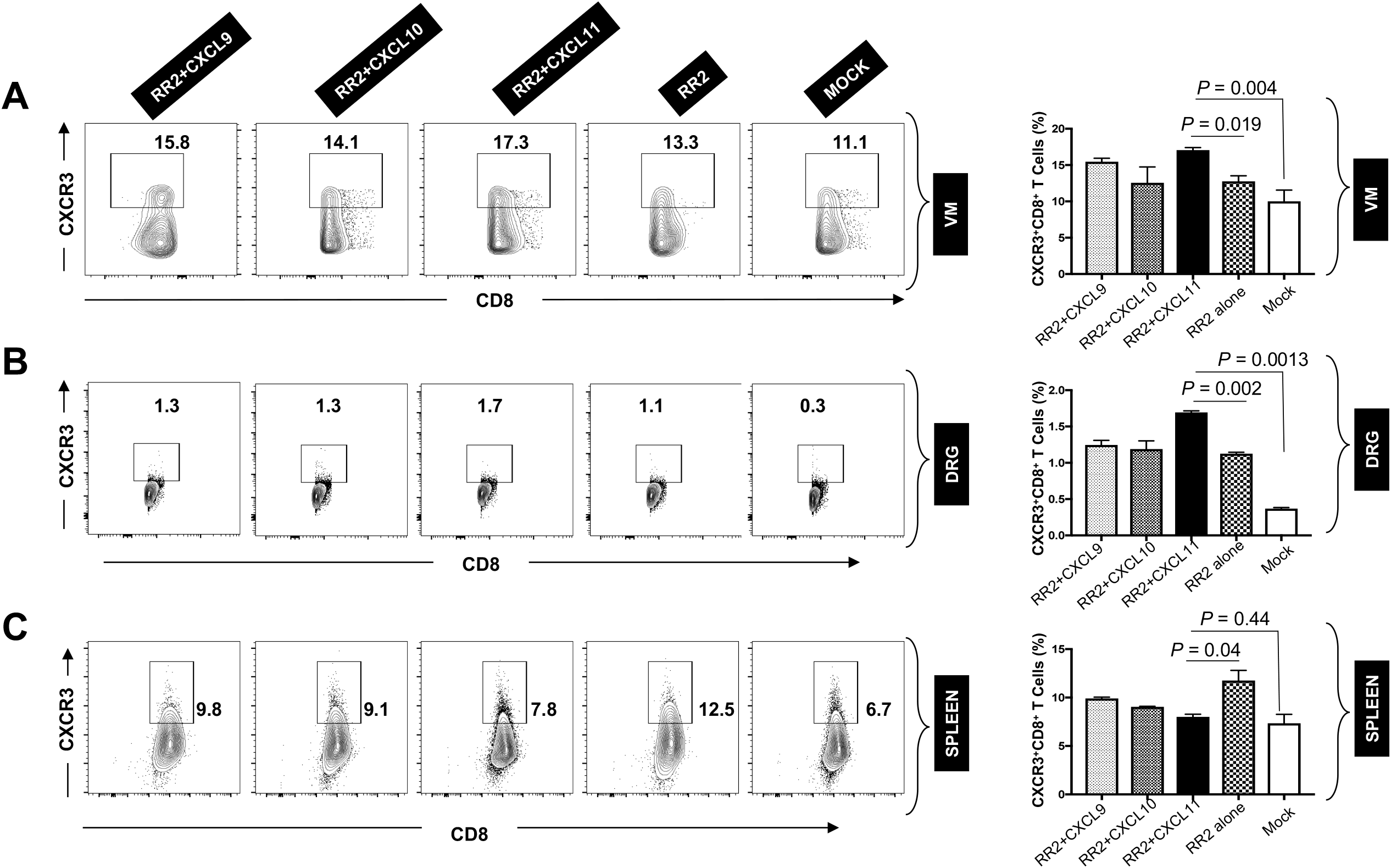
Increased *frequencies of CD8^+^CXCR3^+^ T cells in the vaginal-mucosa and DRG of HSV-2 infected in guinea pigs following therapeutic “prime-pull” vaccination with RR2/CXCL11:* (**A**) Eighty days post-infection, guinea pigs were euthanized, and single-cell suspensions from the spleen (SPL), vaginal mucosa (VM), and DRG were obtained after collagenase treatment. The SPL, VM, and DRG cells were stained for CD8+ and CXCR3 expressing T cells and then analyzed by FACS. Representative FACS data (left panel) and average frequencies (right panel) of CD8^+^ CXCR3^+^ T cells detected in the (**A**) VM, (**B**) DRG (**C**) Spleen of RR2/CXCL9/10/11 and RR2 alone vaccinated and mock vaccinated animals. Cells were analyzed using a BD LSR Fortessa Flow Cytometry system with 4 x 10^5^ events. The indicated P values performed by t-test for significance show statistical significance between prime/pull vaccinated versus RR2 alone and prime/pull vaccinated versus mock-vaccinated control groups.

**5. *Therapeutic prime/pull vaccination of HSV-2 infected guinea pigs with RR2 protein followed by CXCL11 treatment efficiently generate CXCR3 dependent effector memory (T_EM_) in the VM and DRG:*** We next sought to determine the association of various protection parameterswith effector memory in the vaginal mucocutaneous and DRG tissue of HSV-2-infected RR2 + Chemokine and RR2 alone vaccinated guinea pigs. On day 80, after the second and final therapeutic immunization and treatment with chemokines, guinea pigs were euthanized, and single-cell suspensions from the SPL, VM, and DRG tissue were obtained. The effector function of SPL, VM, and DRG-specific CD8 + T cells was analyzed by observing the expression of CD44 on CD8^+^ T-cells by FACS. There was no significant difference observed in the frequencies of T_CM_ in the VM, DRG, and SPL of guinea pigs that were vaccinated with RR2 and treated with chemokines and RR2 protein alone compared to the mock vaccinated group (**Figs. 4A, 4B**, and **4C**). However, significantly higher frequencies of T_EM_ were induced by RR2 vaccinated group treated with chemokines, especially CXCL11, followed by vaccination by RR2 alone compared to those with the mock vaccinated group (i.e., adjuvant alone) in the VM tissues and DRG (**Figs. 4A, 4B** and **4C**).

**Figure 4:**
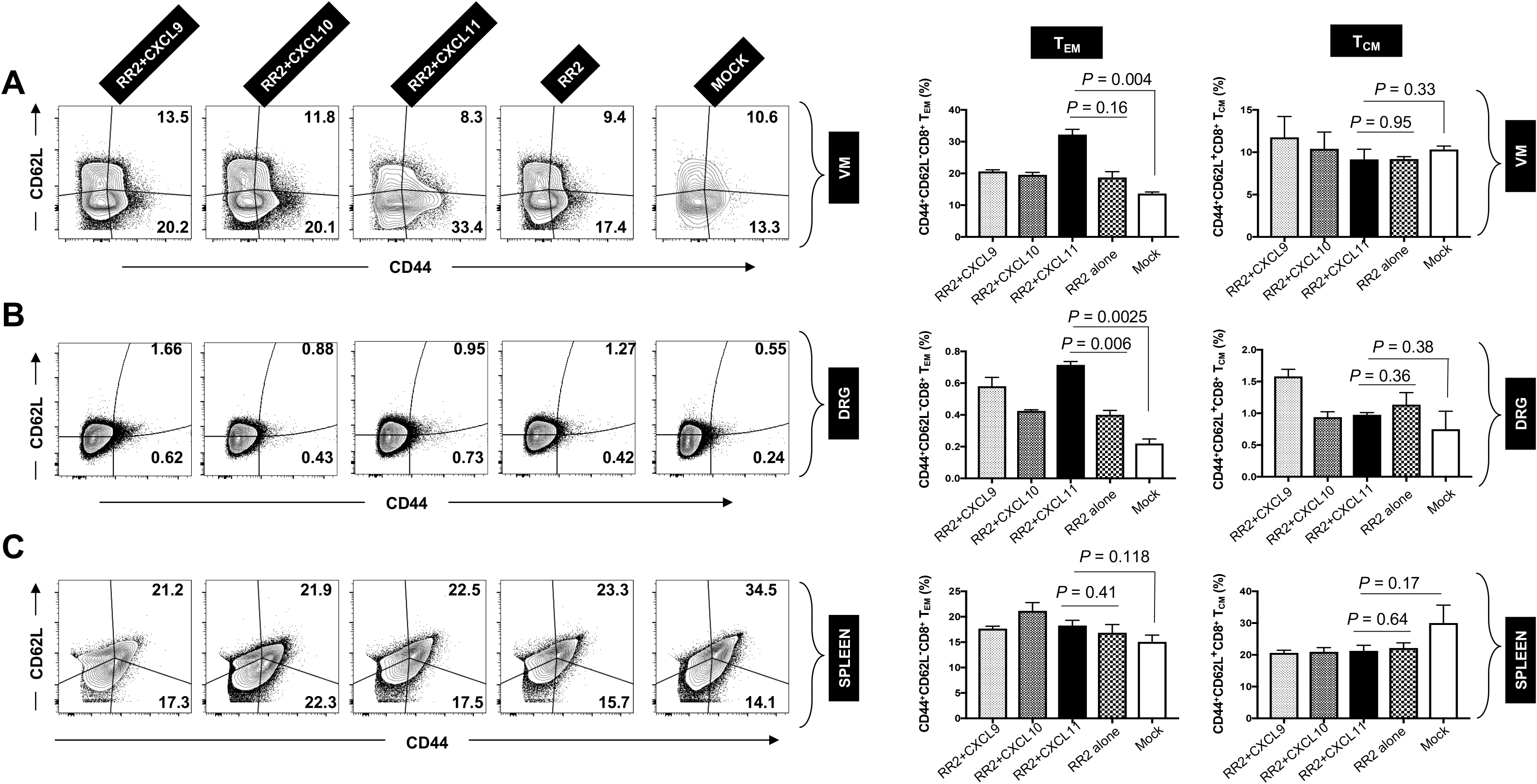
Therapeutic prime/pull vaccination of HSV-2 infected guinea pigs with RR2protein followed by adenovirus expressing chemokine, especially CXCL11 treatment efficiently bolsters effector memory: Eighty days post-infection, guinea pigs were euthanized, and single-cell suspensions from the spleen, vaginal mucosa, and DRG were obtained after collagenase treatment. The SPL, VM, and DRG cells were stained for CD44^+^, and CD62L expressing T cells and then analyzed by FACS. Representative FACS data (left panel) and average frequencies (right panel) of CD44^high^ CD62L^low^ CD8^+^ T_EM_ were detected in the **(A)** VM, (**B**) DRG, **(C)** Spleen of RR2/CXCL9/10/11 and RR2 alone vaccinated and mock vaccinated animals. Higher frequencies of CD44^high^CD62L^low^ CD8^+^ T_EM_ were detected in the VM (**A**) and DRG (**B**) of prime/pull vaccinated guinea pigs, especially the RR2+CXCL11 chemokine treated group. The indicated P values performed by t-test for significance show statistical significance between “prime-pull” vaccinated versus RR2 alone and prime/pull vaccinated versus mock-vaccinated control group.

**6. *Therapeutic prime/pull vaccination of HSV-2 infected guinea pigs with RR2 protein followed by CXCL11 treatment efficiently generate CXCR3 dependent tissue-resident memory (T RM) in the VM and DRG:*** We next determined the association of various protection parameters (i.e., virus shedding and severity and frequency of recurrent genital herpes lesions) with tissue-resident memory that resides at the vaginal mucocutaneous and DRG tissue of HSV-2-infected RR2+Chemokine and RR2 vaccinated guinea pigs. On day 80, after the second and final therapeutic immunization and treatment with chemokine, guinea pigs were euthanized, and single-cell suspensions from the SPL, VM, and DRG tissue were obtained. The effector function of SPL-resident, VM-resident, and DRG-resident CD8^+^T cells was analyzed using both production of CD69 and CD103 expression by FACS. No significant difference was observed in the frequencies of CD69^+^CD103^+^CD8^+^ T-cells in the SPL of guinea pigs vaccinated with RR2 and treated with CXCL11 and RR2 protein alone compared to the mock vaccinated group. However, significantly higher frequencies of CD69^+^CD103^+^CD8^+^ T-cells were induced by RR2 vaccinated group treated with CXCL11, followed by CXCL9, CXCL10, and RR2 alone compared to those with the mock vaccinated group (i.e., adjuvant alone) in the VM and DRG tissues (**Fig. 5A and 5B**).

**Figure 5.**
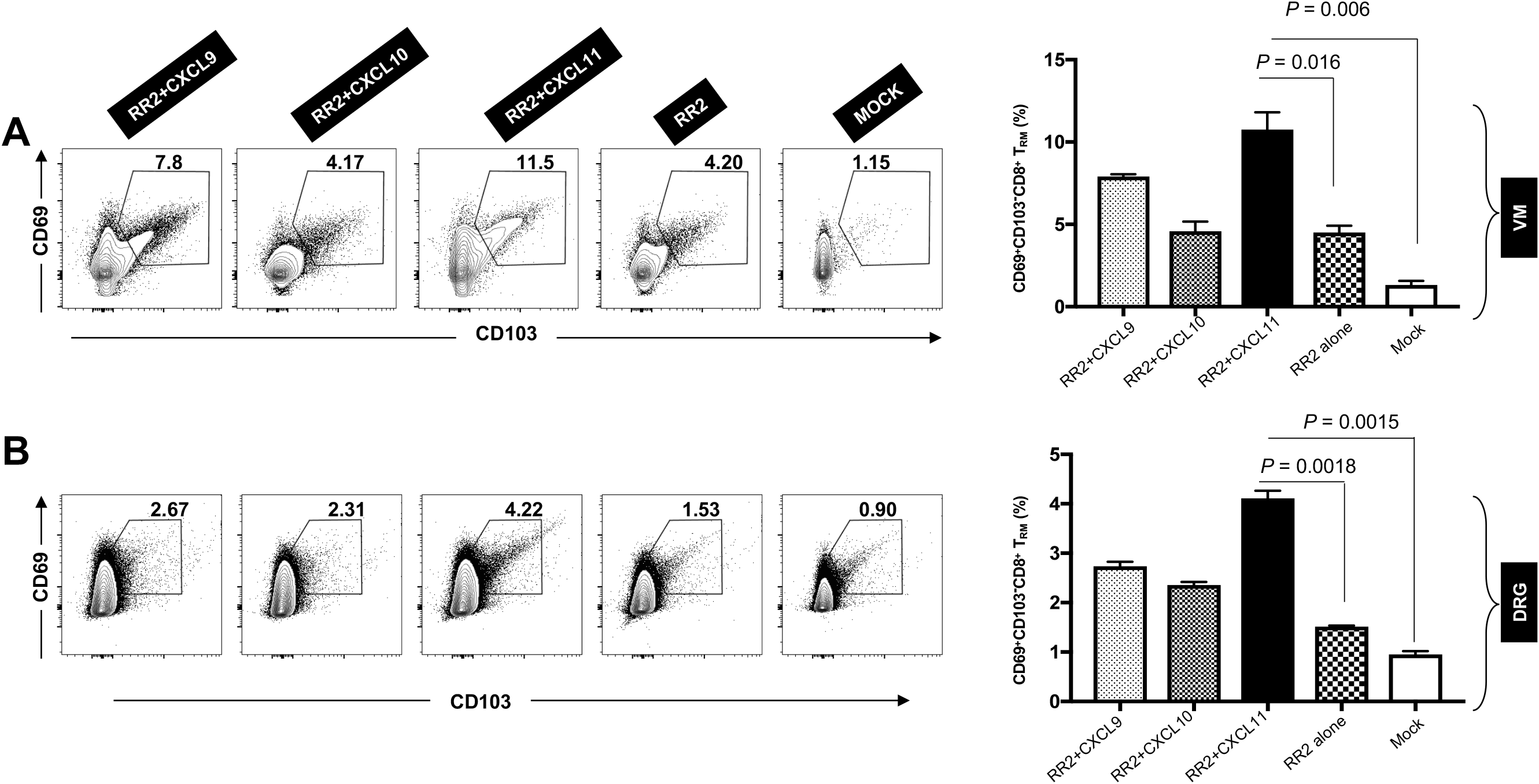
Therapeutic prime/pull vaccination of HSV-2 infected guinea pigs with RR2 protein followed by adenovirus expressing chemokine especially CXCL11 efficiently bolsters tissue-resident memory CD69^+^CD103^+^CD8^+^T_RM_ cells in the VM and DRG of guinea pigs: (**A**) Single-cell suspensions obtained post-infection and immunization from VM, and DRG were stained for memory markers CD69, CD103, and CD8 and analyzed by FACS. Representative FACS data (left panel) and average frequencies (right panel) of CD69^+^CD103^+^CD8^+^ T cells were detected in the (**A**) VM, and (**B**) DRG of RR2/CXCL9/10/11 and RR2 alone vaccinated and mock vaccinated animals. Higher frequencies of CD44^high^CD62L^low^ CD8^+^ T_EM_ were detected in the VM (**A**) and DRG (**B**) of prime/pull vaccinated guinea pigs, especially the RR2^+^CXCL11 chemokine treated group.

**7. *Induced protection from HSV-2 infection following therapeutic prime/pull vaccination with RR2 and treatment with adenovirus containing CXCL11 is associated with more functional tissue-resident IFN-γ^+^CRTAM^+^CD8^+^ T cells:*** We next compared the function of CD8^+^ T cells in the SPL, VM, and DRG of HSV-2 infected guinea pigs following therapeutic prime/pull vaccination with RR2 and treatment with chemokine. On day 80, after the second and final therapeutic immunization and treatment with chemokines, guinea pigs were euthanized, single-cell suspensions from the SPL, VM, and DRG tissue were obtained, and the function of SPL, VM-resident, and DRG-resident CD8^+^T cells were analyzed using both production of IFN-γ and CRTAM expression by FACS. We detected a non-significant difference in the frequencies of IFN-γ-producing CD8^+^T cells in the SPL of guinea pigs that were vaccinated with RR2+Chemokine and RR2 protein compared to the mock vaccinated group (**Figs. 6A** and **6C**). However, significantly high frequencies of IFN-γ producing CD8^+^T cells were observed to be induced by the RR2 vaccinated group treated with chemokine especially CXCL11, followed by the RR2 alone vaccinated group compared to those with the mock vaccinated group in the VM and DRG tissues (**Fig. 6A** and **Fig. 6C**). Similarly, no significant difference was observed in the frequencies of CRTAM expression gated on CD8^+^ T cells in the SPL of guinea pigs vaccinated with RR2 and treated with CXCL11 and RR2 protein compared to the mock vaccinated group (**Fig. 6B** and **Fig. 6D**). However, significantly high frequencies of CRTAM on CD8^+^ T cells were induced by the RR2 vaccinated group treated with CXCL11, followed by CXCL9, CXCl10, and RR2 alone compared to those with the mock vaccinated group in the VM and DRG tissues (**Fig. 6B** and **Fig. 6C**) were detected. In conclusion, these results indicate that therapeutic “Prime-Pull” vaccination of HSV-2-infected guinea pigs with RR2 protein and treated with CXCL11 induced more IFN-γ^+^CRTAM^+^CD8^+^ T cells within the vaginal mucocutaneous tissue and DRG tissue associated with significant protection against recurrent genital herpes.

**Figure 6.**
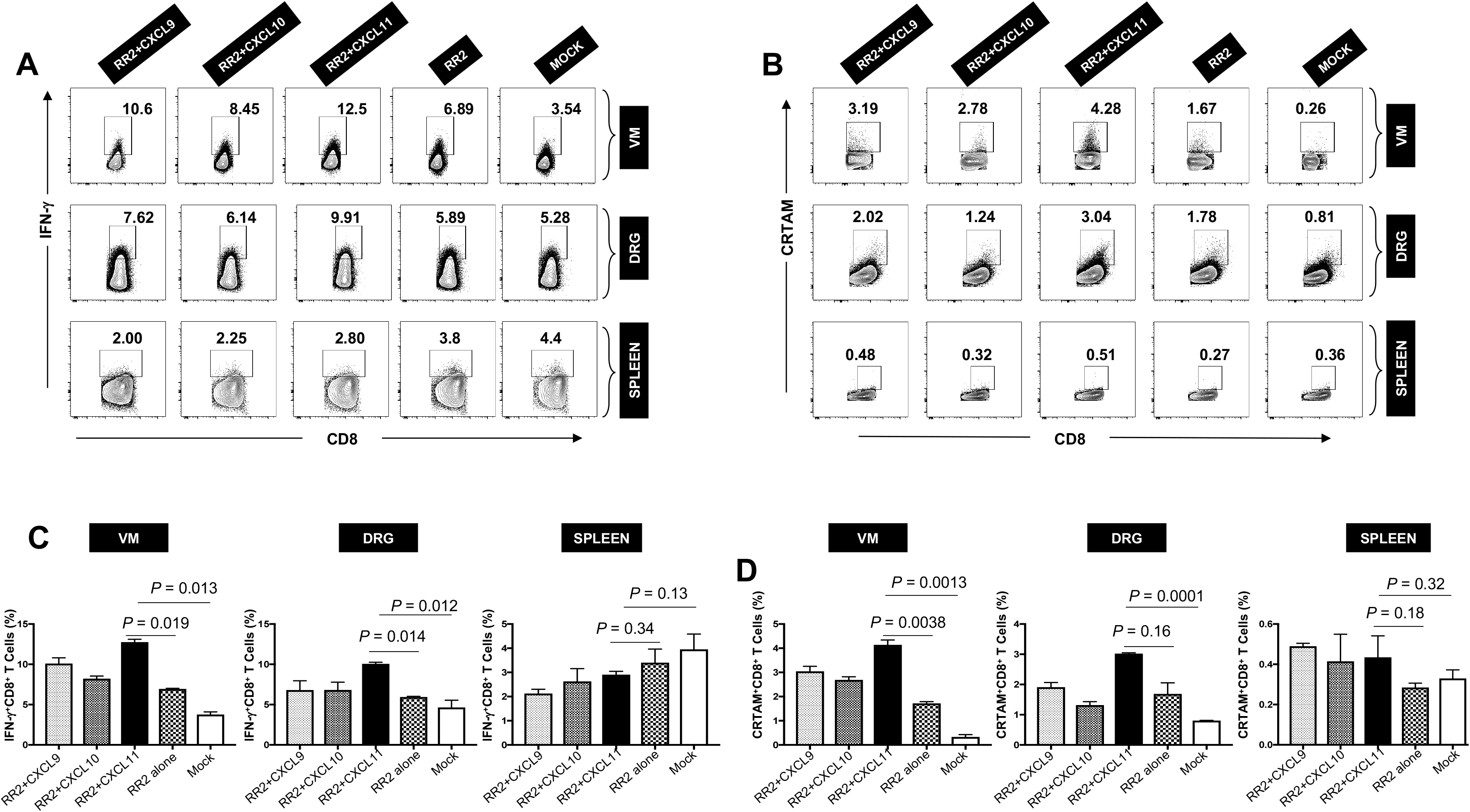
Increased frequencies of functional tissue-resident CD8+ T cells observed in HSV-2 *infected guinea pigs following therapeutic prime/pull vaccination with the RR2 protein/* chemokine especially CXCL11: Functional analysis of CD8^+^ T cells from HSV-2-infected guinea pigs following therapeutic prime/pull vaccination with RR2 protein/chemokine CXCL9/10/11, RR2 alone and mock vaccinated animals. Single-cell suspension from SPL, VM, and DRG were obtained post-infection and immunization. Single cells from SPL, VM, and DRG were stained for functional marker IFN-γ, activation marker CRTAM, and CD8 and analyzed by FACS. Representative FACS data (upper panel) and average frequencies (lower panel) of IFN-γ^+^CD8^+^ T cells were detected in the Spleen, VM, and DRG of RR2+chemokine, RR2 alone vaccinated groups, and the mock vaccinated group. (**A** and **C**) Frequency of IFN-γ^+^CD8^+^T cells per tissue-specific total cells. Representative FACS data (**A**) and average frequencies (**B**) of IFN-γ^+^CD8^+^T were observed in the RR2+chemokine group compared to RR2 alone vaccinated groups and the mock vaccinated group. Similarly, (**B** and **D**) Frequency of CRTAM^+^CD8^+^T cells per tissue-specific total cells. Representative FACS data (**B**) and average frequencies (**D**) of CRTAM^+^CD8^+^T were observed in the RR2+chemokine group comparison compared to RR2 alone vaccinated groups and the mock vaccinated group.

## DISCUSSION

The morbidity and the socioeconomic burden related to recurrent genital herpes highlight the need for a therapeutic herpes vaccine that can mitigate the impact of the disease. Despite its toll, herpes has generally been seen as a “marginal” disease. At present, only a handful of pharmaceutical companies and academic institutions have invested in herpes vaccine research over the past few decades (27-29). The past clinical trials that used glycoproteins gB and gD as the primary antigen in herpes vaccines met failure. Although these glycoproteins are commonly targeted by the immune system, they did not provide sufficient protection against recurrent herpetic disease when included in the vaccine (30-35). In the present study, lessons learned from the failed gB/gD-based vaccines led us to hypothesize therapeutic subunit vaccine containing non-gB/gD “asymptomatic” HSV-2 proteins that can selectively induce CD4^+^ and CD8^+^ T_RM_ cells from naturally “protected” asymptomatic women would decrease the frequency and severity of the recurrent herpetic disease.

Recurrent genital herpetic disease is one of the most common sexually transmitted diseases, with a worldwide prevalence of infection predicted to be over 3.5 billion individuals for HSV-1 and over 536 million for HSV-2 (32-33). However, given the staggering number of individuals already infected with HSV-1 and HSV-2, there is a significant unmet medical need for a therapeutic herpes vaccine to reduce viral shedding and alleviate herpetic disease in symptomatic patients. Historically, it has been harder to develop therapeutic vaccines because the virus-HSV-1 and HSV-2 employ many strategies to evade the host immune system, which are in place in already infected individuals (36-39). Thus, scientists relied on developing prophylactic vaccines to prevent new infections in seronegative individuals. Moreover, the potential for recombination between HSV-1 and HSV-2 has given rise to HSV recombinant strains, that are widely circulating (40). The majority of HSV-1 and HSV-2-seropositive individuals are asymptomatic, implying that such individuals can readily and quietly transmit the virus to their partners. Global herpes prevalence stresses the urgency of developing a therapeutic vaccine. In our previous research, the protective efficacy of eight HSV-2 proteins that were highly recognized by the immune system from naturally protected asymptomatic individuals as therapeutic vaccine candidates were compared. Among others, we found that immunization using HSV-2 RR2 protein demonstrated significant protection against recurrent genital herpes disease. Moreover, RR2 protein-induced HSV-2 specific antibody and T-cell responses correlated with reduced viral shedding and disease severity (26). In addition, previous research has identified a CD8^+^ T cell population located near nerve endings that control reactivated HSV (41-42). This suggests that the presence and activity of CD8^+^ T cells in the mucosa are critical for controlling HSV reactivation and subsequent disease outcomes.

The HSV-2 ribonucleotide reductase (RR2) is a major target of HSV-2-specific CD8 T cells in humans. It consists of two heterologous protein subunits. The small subunit (RR2) is a 38-kDa protein encoded by the UL40 gene, and the large subunit (RR1), designated ICP10, is a 140-kDa protein encoded by the UL39 gene (43). RR2 protein can boost neutralizing antibodies and increase the numbers of functional IFN-γ^+^CRTAM^+^ CD8 T cells within the VM tissues. While systemic memory T cells can migrate freely through organs such as the spleen and liver, others such as the intestines, lung airways, central nervous system, skin, and vagina, are restrictive for memory T cell entry (44). In the latter tissues, inflammation or infection is often required to permit entry of circulating activated T cells to establish a tissue-resident memory T cell pool that composes a separate compartment from the circulating pool (45). Given that the occurrence of inflammation in the reproductive tissue may preclude the infection or reactivation of the virus, we investigated an alternative approach to recruit virus-specific T cells into the vaginal mucosa without inducing local inflammation or infection i.e. using chemokines. This strategy referred to as the “Prime-Pull” strategy, involves priming the immune system with an initial vaccination and then “pulling” the immune response to the site of infection using specific chemokines. The goal is to increase the number of HSV-specific CD8^+^ T cells that can effectively control reactivated HSV and reduce recurrent disease and viral shedding. Chemokines are naturally produced by our immune system and could serve as safer and more reliable molecules to attract immune cells (46). The CXC chemokine ligand 10 (CXCL10)/CXC chemokine receptor 3 (CXCR3) pathways are critical in promoting T cell immunity against many viral infections. A “Prime-Pull” Therapeutic Vaccine can boost Neutralizing IgG/IgA antibodies and boost the number and function of antiviral CD4^+^ and CD8^+^ TRM cells within the cervical genital mucocutaneous (CGMC) and DRG tissues and thus expected to help stop the virus reactivation from latently infected DRG, virus shedding, and virus replication in CGMC, thus curing or reducing recurrent genital herpes disease. In one of the “Prime-Pull” strategies, a topical chemokine was applied to the genital mucosa after subcutaneous vaccination to pull HSV-specific CD8^+^T cells that were found to be associated with decreased disease upon challenge with HSV-2 (47-49). The chemokine/CXCR3 pathway also affects TG- and cornea-resident CD8^+^ T cell responses to recurrent ocular herpes virus infection and disease (50-52). Chemokines can also be co-delivered in a DNA vaccine for immunomodulation. Pre-clinical studies in HSV have shown immuno-potentiation of DNA vaccines by co-delivery of chemokines such as CCR7 ligands and IL-8, RANTES delivered to the mucosa (53). In this study, we establish the therapeutic efficacy of the “Prime-Pull” strategy using HSV-2 protein for immunization (Prime) and a chemokine (Pull) to increase the number of HSV-2-specific T-cells in the vaginal mucosa of guinea pigs. The localized increase in T cells frequency was attained by using adenovirus expressing guinea pig-specific mucosal chemokine CXCL9/10/11.

Guinea pigs were immunized (Primed) using previously studied RR2 protein and RR2-specific T-cells were pulled in VM using CXCL9/10/11. We hypothesized that the chemokine-based pulling would increase the functional, tissue-resident, IFN-γ-producing T cells in the VM. Our studies demonstrate that the chemokines especially CXCL11 treatment could pull the RR2-specific CD4^+^ and CD8^+^ T-cells in the VM. The chemokine ligand 11, also known as IFN-inducible T cell α chemoattractant (I-TAC), mediates the recruitment of T cells, natural killer (NK) cells, and monocytes/macrophages at sites of infection, predominantly through the cognate G-protein coupled receptor CXCR3 (54). This signaling axis has been implicated in several physiological activities, including immune cell migration, differentiation, and activation (55). In addition, the affinity of CXCL11 for CXCR3 is the highest among the three selective ligands i.e. CXCL9, CXCL10, and CXCL11 (53-56). We show an increase in the number and function of tissue-resident memory CD8^+^ T cells in the vaginal mucosa of vaccinated guinea pigs treated with CXCL11 than that observed with other chemokines and with RR2 alone. The pulling of T-cells manifested in a significant reduction of virus shedding and decreased recurrent genital herpes lesions. The viral reduction was significantly higher in vaccinated guinea pigs treated with CXCL11 than that observed with other chemokines and with RR2 alone. Upon evalution of the expression of CXCR3 on CD8^+^ T cells in the VM, DRG, and spleen of guinea pigs, we observed an increased number and percentage of CXCR3^+^CD8^+^ T-cells in the VM and DRG of guinea pigs treated with chemokine/RR2 compared to RR2 alone. The reduced frequency of CXCR3^+^CD8^+^ T cells in mock and RR2 alone highlights the role of chemokine/RR2 in the migration of CD8^+^ T cells in the VM and DRG of chemokine-treated guinea pigs. Moreover, the expression of CXCR3 on CD8^+^ T cells was predominant in the DRG of vaccinated guinea pigs treated with CXCL11. We believe that CXCL11-dependent therapy may be a potential approach for viral infections.

In conclusion, our study demonstrates that immunizing guinea pigs with an immunogenic HSV-2 protein RR2, and treatment with adenovirus-expressing chemokine provides better protection against recurrent genital Herpes. This strategy seems to prevent the migration of HSV-2 from mucosa to neurons leading to decreased reactivation and viral shedding. We hypothesize that CD8^+^ T cells reduce neuronal infection or viral replication within neurons. In addition, the presence and increase in activated CD8^+^ T cells in genital mucosa also suggest that viral establishment and replication may be inhibited at the entry site. However, the exact mechanism by which the T cells function or control the infection is not yet established. The presence and magnitude of appropriate chemokines at the site of infection are important to achieve maximum protection and may be achieved by optimizing “Prime-

Pull” for 100% efficacy. This type of immunotherapy can help recruit and establish resident T cells that can provide protection not only against genital herpes but also other types of sexually transmitted diseases.

## ACKNOWLEDGEMENTS

This work is dedicated to the memory of the late Professor Steven L. Wechsler “Steve” (1948-2016), whose numerous pioneering works on herpes infection and immunity laid the foundation for this line of research.

